# Neural and linguistic differences explain priming and interference during naming

**DOI:** 10.1101/547745

**Authors:** Tao Wei, Tatiana T. Schnur

**Affiliations:** College of Psychology and Sociology, Shenzhen University, Shenzhen, Guangdong 518061,China; Baylor College of Medicine, Department of Neurosurgery, Houston, TX 77030, USA

**Keywords:** language production, fMRI, priming, interference, functional connectivity

## Abstract

When naming an object, humans are faster to produce the name (“cat”) if immediately having named a related object (“dog”) but paradoxically slower to name the same object (“cat”) if there are intervening speech acts (Wei and Schnur 2019). This dependence of behavior on prior experience is ubiquitous in other domains, often termed “priming” (if behavior is speeded) or “interference” (if behavior is slower). However, it is unknown the changes in the language system (conceptual, lexical, and/or connections between representations) and corresponding brain mechanisms which create these paradoxical effects on the same speech act. Using fMRI during simple picture naming, we observed distinct brain regions and different connections associated with priming and interference. Greater priming was associated with increased activation in the ventral occipitotemporal cortex, while greater interference was associated with decreased functional connectivity between the left posterior temporal and angular gyri. To provide neural evidence of where in the language system priming and interference in naming occur, we assayed the response of different brain areas to conceptual or lexical aspects of speech. The brain and language systems adapt to prior naming experience by modulating conceptual representations during priming, but modulating conceptual, lexical and the mapping between representations during interference.

## Introduction

Speech is complex, requiring the retrieval of different types of information for a successful speech act. For example, to successfully communicate a thought, a speaker needs to match the idea of what she wants to convey to a specific concept (e.g., ramen), activate the conceptual representations, which then need to match a lexical (word) form in the language she speaks, and then activate the sounds and motor programs that are associated with the word she wishes to say. The question we addressed here is how the brain and language systems are shaped by previous speech acts which then impact future speech.

What we name in the past affects word production in the future. For instance, naming a target picture (e.g., CAT) is speeded up when it is preceded by a related prime picture/word (e.g., DOG) compared to an unrelated prime (e.g., VASE; Lupker 1988; Wei and Schnur 2019). Intriguingly, future word production is slowed down in almost identical circumstances (e.g., Brown 1981; Wheeldon and Monsell 1994; Damian et al. 2001; Howard et al. 2006; Schnur et al. 2006; Vitkovitch et al. 2006; Belke 2013). Recently, using a simple within-subject and within-item single picture naming task we demonstrated that the emergence of priming or interference in word production depends on the interval between two naming occurrences (Wei and Schnur 2019). To address the neural and cognitive changes responsible for this paradoxical naming phenomenon, in the current study we combined task functional and connectivity magnetic resonance imaging (MRI) with a linguistic assay approach. To our knowledge, this study is the first to identify within the same participants, items, and task how the language system and supporting brain mechanisms change to produce faster or slower speech.

In cognitive models of language production, priming and interference are generally assumed to occur at different levels of processing during word production (e.g., Dell 1986; Caramazza 1997; Levelt et al. 1999). That producing a word is sped up or slowed down suggests that different processes during word production are affected by prior naming experience. Just like priming widely reported in other language tasks is hypothesized to occur during access to word meanings (word reading, e.g., Meyer et al. 1975; lexical decision (a decision as to whether a string of letters is a word or not), Meyer and Schvaneveldt 1971; conceptual decision (judge whether the word referred to is a concrete object), e.g., McRae and Boisvert 1998; see Neely 1991 for review), the priming effect in word production has been attributed to spreading activation at the conceptual level of processing. When producing the name of a target picture, a previously produced semantically related word (the prime) will speed up target naming compared with unrelated primes because semantically related primes share more conceptual information with targets and thus send greater spreading activation to targets, which facilitates subsequent target naming (e.g., Sperber et al. 1979; Huttenlocher and Kubicek 1983; Lupker 1988). In contrast, to account for interference in naming, it has been assumed that semantically related vs. unrelated primes evoke greater competitive selection (Howard et al. 2006; Roelofs 2018) or decreased activation (Oppenheim et al. 2010) at the lexical level of processing, which hampers subsequent target naming.

While priming and interference are thought to involve different cognitive processes (conceptual vs. lexical), similar brain regions are associated with both effects. Although no study to our knowledge has identified the neural locus of priming in naming, numerous fMRI studies have investigated the neural basis underlying priming in lexical decision (e.g., Rossell et al. 2003; Gold et al. 2006; Wible et al. 2006) and word reading (e.g., Wheatley et al. 2005). During lexical decision and word reading tasks, semantically related vs. unrelated primes modulated activation in distributed brain regions, including the bilateral ventral occipitotemporal cortex (vOTC), left middle and superior temporal gyri, and precuneus cortex (Gold et al. 2006; Rossell et al. 2001; 2003; Wible et al. 2006). Interestingly, similar regions are active during interference in naming. During naming paradigms where interference was observed behaviorally (e.g., Howard et al. 2006; Schnur et al. 2006; Wheeldon and Monsell, 1994), activation in a range of cortical regions, including the left middle and inferior temporal gyri and the left inferior frontal gyrus, was affected by whether a semantically related picture was naming previously (de Zubicaray et al. 2006; 2013 2014; 2015; Schnur et al. 2009). The involvement of the left middle temporal gyrus has been interpreted as evidence consistent with a lexical locus of interference (de Zubicaray et al. 2006; 2013), because this region is reported elsewhere as involved in lexical processing during speech production (see Indefrey and Levelt 2004 for review). However, similar regions have also been found to be involved in conceptual processing (see Martin 2007 and Binder and Desai 2011 for review). Without directly testing the cognitive function (conceptual vs. lexical) of the regions sensitive to the manipulation of prior naming experience, we cannot infer where in the language system priming and interference happen, solely based on their neural loci. Moreover, because no studies have yet examined the neural loci of priming in naming and compared it with the loci of interference in naming, we do not know whether the same brain region is modulated by prior naming experience and induces both effects via different mechanisms depending on the recency of naming experience or whether the paradoxical effects occur at different but nearby regions. Therefore, to understand how the brain and language systems are shaped by prior naming experience, we investigated priming and interference together and assessed the cognitive roles of corresponding brain regions directly.

In addition, beyond modulating regional brain activation, prior naming experience may also change the functional connectivity patterns between regions, highlighting another potential mechanism underlying the paradoxical effects in word production. For example, Ulrich and colleagues (2014) showed that in a lexical decision task, the masked (subliminal) semantically related vs. unrelated primes differentially modulated the activation of vOTC as well as the functional connectivity associated with vOTC. Further, the difference of functional connectivity strength between the related and unrelated condition was significantly correlated with the magnitude of the priming effect across participants. This suggests that previously processed stimuli (the primes) changed the coupling between brain regions, which affected the efficiency of mapping between information stored in different regions and in turn influenced participants’ behavior. Indeed, some computational models (Howard et al. 2006; Oppenheim et al. 2010) assume that prior naming experience modulates the mapping between conceptual and lexical representations, directly affecting the lexical level of processing resulting in interference in naming. Thus, to comprehensively examine how the brain is affected by prior naming experience, we tested whether naming experience modulates both functional connectivity patterns and regional brain activity.

In the current fMRI study, we examined the mechanisms underlying the paradoxical effects of priming and interference in word production with the following manipulations. First, we measured fMRI BOLD response when participants named pictures in a naming paradigm which induces both priming and interference (i.e., the priming-interference picture naming task; Wei and Schnur 2019). When a prime and target picture are named temporally adjacently (lag0), the semantically related naming experience primes subsequent naming. When a prime and target picture are separated temporally by two unrelated pictures (lag2), the semantically related naming experience hampers naming performance. Because this task creates priming or interference during naming of the same item within the same participant, this enabled us to directly compare the neural mechanisms underlying both effects without confounds which could explain potential differences between effects (e.g., differences between tasks or participants). Second, in addition to brain activation, we examined whether prior naming experience modulates functional connectivity between brain regions and thus gives rise to priming or interference. Third, to understand the relationship between the neural and behavioral changes caused by prior naming experience, using correlational analyses we tested whether the neural changes in terms of activation and functional connectivity were directly related to the individual differences in behavioral changes (response times) caused by prior naming experience. Fourth, we included an independent naming task designed to assay the response of different brain areas to conceptual vs. lexical aspects of language processing (i.e. a linguistic assay task). In psycholinguistic studies, concept familiarity (how familiar a concept is in a person’s individual experience) and lexical frequency (how often a word occurs in text or speech) are commonly used to index conceptual (e.g., Hirsh and Funnell 1995; Funnell and De Mornay Davies 1996; Lambon Ralph et al. 1998; Rogers et al. 2015; Woollams et al. 2008) and lexical processing (e.g., McClelland and Rumelhart 1981; Dell 1986; Vitkovitch and Humphreys 1991; Caramazza 1997; Rastle and Coltheart 1999; Finocchiaro and Caramazza 2006; Strijkers et al. 2010; cf. Levelt et al. 1999) respectively. Employing a parametric fMRI approach, Graves et al. (2007) and Wilson et al. (2009) identified different brain regions involved in conceptual and lexical processing during naming by testing a region’s activation sensitivity to concept familiarity or lexical frequency. Following the same logic, to identify the cognitive role (conceptual vs. lexical) of brain regions modulated by prior naming experience, we tested whether the brain regions modulated by prior naming experience overlapped with brain regions most responsive to conceptual vs. lexical features.

## Materials and Methods

### Participants

30 right-handed native English speakers from Rice University participated in this study. They were safety-screened and reimbursed in accordance with the Institutional Review Board at Rice University. Data from four participants were excluded: one due to experimenter error, two due to the interruption caused by a noise cancellation failure of the headphones, and one due to head motion over 2 mm, resulting in a total of 26 participants (18-22 years old).

### Materials, Design and Procedure

#### Priming-Interference Picture Naming Task

To uncover the neural basis of the paradoxical priming and interference effects caused by prior naming experience, we adopted a simple picture naming task (referred to as the priming-interference naming task; Wei and Schnur 2019) where participants name pictures one at a time. We selected this task because of its simplicity and its robustness in creating within-subject priming and interference during the naming of the same items, across different languages and groups of participants. In the priming-interference naming task, participants are either faster to name a target picture (priming) or slower to name the same target (interference) depending on the recency of naming a semantically related vs. unrelated previous picture. Specifically, we observed priming and interference during naming of the same items by manipulating the number of unrelated intervening trials (lag0 vs. lag2) between two semantically related vs. unrelated (relatedness) naming occurrences. Following Wei and Schnur (Experiment 2 2019), we manipulated relatedness (related vs. unrelated) and lag (lag0 vs. lag2) as within-subject and within-item factors which reduces the variance associated with individual differences and different materials. Stimuli included 52 prime, 52 target and 104 filler pictures from the Bank of Standardized Stimuli (BOSS, Brodeur et al. 2014) and online resources. Each trial consisted of four pictures: a prime, a target and two filler pictures (Figure 1). In the related condition, the prime and target in a pair were semantically related to each other. To create the unrelated condition, the same stimuli were regrouped so that the prime and target in a pair were unrelated to each other. In the lag0 condition, two fillers were presented before the prime, and the prime and target were presented adjacently (i.e., filler, filler, prime, target). In the lag2 condition, two fillers were inserted between the prime and target (i.e., prime, filler, filler, target).

**Figure 1.**
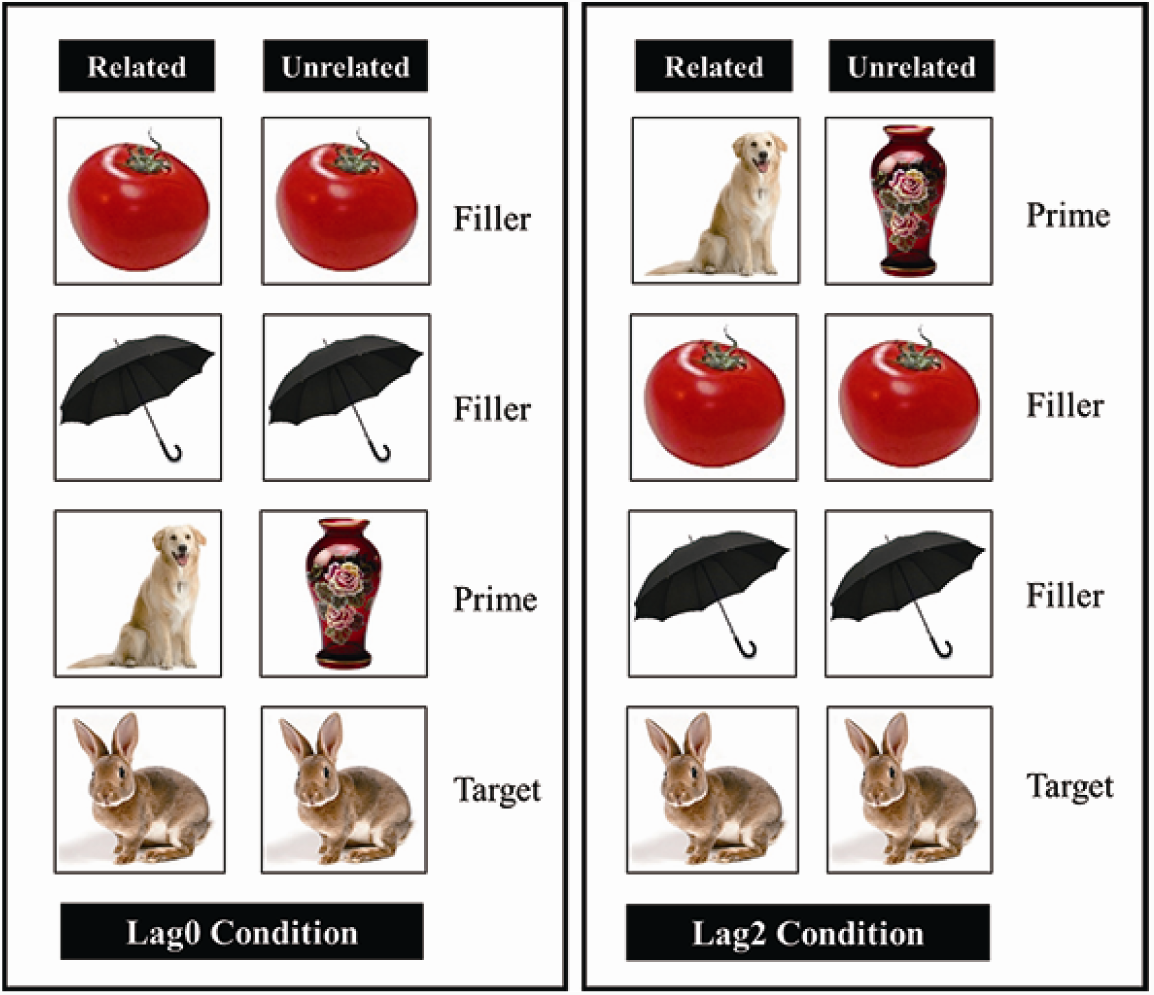
Example of stimuli from a trial across different conditions: relatedness (related vs. unrelated) × lag (lag0 vs. lag2).

During the experiment, we instructed participants to name pictures as quickly and accurately as possible. Each participant named the same picture four times, once in each condition (related × lag). This resulted in a total of 208 trials (832 pictures) and four runs of 52 trials per participant. In the same run, no picture appeared twice and there were 13 trials from each of the four conditions (relatedness × lag). For each picture presented within a trial, we presented a 500 ms fixation point (+) followed by a picture stimulus presented for 1000 ms, followed by a 500 ms blank screen, resulting in a duration of 8 sec. for each trial of four pictures. The inter-trial interval (ITI) was jittered to be 0, 2, or 4 sec. (average = 2 sec.) during the display of fixation point (+). We included the ITI to promote stimulus jittering and serve as a baseline in the fMRI data analyses (see Data Analysis). The pictures from different conditions and jitters were presented in a randomized order, optimized using make_random_timing.py and 3dDeconvolve functions in AFNI. The naming task took approximately 38 minutes (including three 1-minute inter-run-intervals).

#### Linguistic Assay Naming Task

We designed the linguistic assay naming task to identify the cognitive role (conceptual vs. lexical; cf. Graves et al. 2007; Wilson et al. 2009) of the brain regions associated with the paradoxical effects created by prior naming experience. The stimuli comprised 142 photographs from the BOSS set (Brodeur et al. 2014) and 24 photographs from online resources. Items did not overlap with the priming-interference naming task stimuli. We selected the stimuli based on variability in both word frequency and concept familiarity. We determined picture stimuli word frequency using The English Lexicon Project (Balota et al. 2007) with log10 transformation (mean: 3.49, range: 1.85-5.27). We determined picture stimuli concept familiarity from norms provided in the BOSS stimuli set (Brodeur et al. 2014). Considering that 24 picture stimuli in the linguistic assay task were not from the BOSS stimuli set, participants performed a concept familiarity rating task on all materials used in the linguistic assay task three days before their fMRI experiment using an on-line survey tool (Qualtrics, https://riceuniversity.co1.qualtrics.com). Following Fiez and Tranel (1997), we asked participants to rate the level to which they were familiar with the objects on a 5-point scale (1, very unfamiliar and 5, very familiar). We instructed participants to rate the concept itself rather than the picture of the object and encouraged them to employ the full range of scale options throughout the set of pictures. For the 142 pictures from the BOSS stimuli set, the concept familiarity ratings from our participants (mean: 4.15, range: 2.69-5) were highly correlated with the concept familiarity ratings from the BOSS norms (*r* =.88, *p* <.001,), and were not correlated with the lexical frequency of picture names (*r*=.07, *p*=.38).

Participants followed the same instructions from the priming-interference naming task, to name pictures as quickly and accurately as possible. Each trial lasted 2 sec.: a fixation point (+) was displayed for 500 ms, followed by the picture stimulus for 1000 ms and a blank screen for 500 ms. The ITI with a display of fixation point (+) was jittered at 0, 2, or 4 sec. (average = 2 sec.) to promote stimulus jittering and serve as a baseline for the fMRI data analyses (see Data Analysis). All trials were divided equally into two runs per participant. The pictures and jitters were presented in a randomized order. The linguistic assay task took 12 minutes with a 1-minute break between two runs and always occurred immediately following the priming-interference task.

### Data Acquisition

#### Behavioral Data

We used the E-Prime 2.0 software (Psychology Software Tools, Pittsburgh, PA) to program and run the picture naming experiment inside the scanner. The visual display (i.e., instruction and picture stimuli) was presented on an LCD panel and back-projected onto a screen positioned at the front of the magnet. Participants lied down in the scanner and viewed the display on a mirror positioned above them. The OptoActive active noise cancelling system was used to reduce the noise during scanning and record the vocal responses.

#### Imaging Data

MRI scanning was performed on a 3T Siemens Trio MRI scanner at the Baylor College of Medicine Core for Advanced Magnetic Resonance Imaging (CAMRI). All participants first underwent one structural scan, followed by four functional runs of priming-interference naming and two functional runs of the linguistic assay task. The high-resolution 3-dimension structural image was acquired with MPRAGE sequence in the axial plane (repetition time (TR) = 2600 ms, echo time (TE) = 3.03 ms, flip angle (FA) = 8°, matrix size = 256 × 256, voxel size = 1 × 1 × 1 mm^3^) for normalization (see below). Functional data were collected in the Echo-Planar Imaging (EPI) sequence as follows: TR = 2000 ms, TE = 30 ms, FA = 72°, matrix size = 100 × 100, FOV = 160 mm, voxel size = 2 × 2 × 2 mm^3^. For each volume, 62 2-mm axial slices were collected to cover the participants’ entire brain.

### Data Analysis

#### 1. Behavioral Data

To score responses for accuracy and obtain response times, we first used the noise reduction function in Audacity (www.audacityteam.org) to remove noise due to scanner activation while leaving the speech signal intact. A native English speaker transcribed the sound files, extracted the RTs using CheckVocal (Protopapas 2007) and encoded responses as correct (matched the target) or incorrect (wrong name for the target, sound error, or omission). For the RT analyses we used RTs from correct responses. For the error analyses, we analyzed target wrong names and omissions. To test for the priming and interference effects caused by past naming experience based on the RT data of the priming-interference picture naming task, we conducted planned paired *t*-tests in lag0 and lag2 respectively, including subjects (*t*_1_) and items (*t*_2_) as random factors, and relatedness as a fixed within-subject and within-item factor. For the linguistic assay naming task, to confirm the influence of lexical frequency and concept familiarity on picture naming, we averaged response times/accuracies across participants for each picture and then performed Pearson correlations between mean response times/accuracies with concept familiarity ratings and log-transformed lexical frequencies of picture names.

#### 2. Imaging Data Preprocessing

Preprocessing of imaging data was identical for both priming-interference and linguistic assay naming tasks using the software package Statistical Parametric Mapping (SPM12, http://www.fil.ion.ucl.ac.uk/spm/software/spm12/). The first three EPI volumes (6 sec.) of each functional run were discarded to allow for signal equilibration. After slice timing correction, the EPIs were realigned to the session mean. To wrap participants’ EPI images to the Montreal Neurological Institute (MNI) space, each participant’s T1 image was first co-registered to the mean EPI. Then, the co-registered T1 image was segmented into gray and white matter, to create a template using the DARTEL toolbox (Ashburner 2007) in SPM12. The resulting individual deformation field was applied to the T1 and EPI images for spatial normalization to MNI space. The normalized EPI images were written to 2 × 2 × 2 mm and smoothed with a 6 mm full-width at half-maximum isotropic Gaussian kernel to decrease spatial noise prior to statistical analysis.

#### 3. Whole-Brain Univariate Activation Analysis

We conducted whole-brain univariate activation analyses to identify the brain regions whose activation was affected by prior naming experience (relatedness X lag). After preprocessing the priming-interference picture naming imaging data, for each participant we used a general linear model (GLM) to model the fMRI BOLD response to each picture. To account for voxels whose activation differed depending on the target picture condition (i.e. conceptual relatedness X lag), four predictors were formed with the onsets of target picture: lag0-related, lag0-unrelated, lag2-related and lag2-unrelated. Furthermore, two predictors representing the onsets of primes and fillers and six head motion parameters as nuisance covariates were added to the design matrix. Then all predictors were convolved with the canonical hemodynamic response function (HRF). To remove low-frequency scanner drifts, data were high-pass filtered with a frequency cutoff at 128 s. In the first level of the GLM, contrast images were computed on each participant to represent the main effects of lag0-related, lag0-unrelated, lag2-related, and lag2-unrelated trials versus baseline (jitter trials). In the second level of the GLM, to identify the brain regions underlying priming and interference naming effects respectively, across participants we performed *t*-contrasts between lag0-related and lag0-unrelated images and between lag2-related and lag2-unrelated images. Significant clusters of activation were determined at a voxel height threshold of *p* < 0.001 and a family-wise error (FWE) -corrected cluster threshold of *p* < 0.05. Lastly, we performed a conjunction analysis (Nichols et al. 2005) to test whether priming and interference involve the same regions.

#### 4. Functional Connectivity Analysis

To identify how prior naming experience (relatedness X lag) modulates the functional connectivity between brain regions during naming, we performed functional connectivity analyses using the generalized form of context-dependent psychophysiological interactions approach (gPPI, McLaren, Ries, Xu, and Johnson 2012; http://www.nitrc.org/projects/gppi). Specifically, we examined the relatedness-modulated functional connectivity of the brain regions identified by the whole-brain univariate activation analysis for the priming (lag0) and interference (lag2) conditions respectively. First, the regions identified in the voxel-wise analysis were defined as seed regions. Then, the time series from each seed region was first deconvolved to represent neuronal activation and then multiplied with the design matrix used in the univariate analysis to form an interaction term. This interaction term was further convolved with the HRF to form the gPPI regressor. The resulting time series was regressed against all other voxels in the brain to form the effective functional connectivity map of the corresponding seed region.

We were interested in how the functional connectivity pattern associated with a seed region was modulated by prior naming experience (relatedness X lag). To explore the functional connectivity pattern underlying priming, we performed *t*-contrasts between lag0-related and lag0-unrelated maps for the priming seed regions. Similarly, we performed t-contrasts between lag2-related and lag2-unrelated maps for the interference seed regions to uncover the functional connectivity pattern underlying interference. Significant clusters of functional connectivity were determined at a voxel threshold of *p* < 0.001 and a FWE-corrected cluster threshold of *p* < 0.05.

#### 5. Linguistic Assay Parametric Analysis

To determine the cognitive role (conceptual vs. lexical) of brain regions and functional connectivity associated with priming and interference, we conducted the following ROI-based parametric analysis. First, we defined the clusters found in the whole-brain univariate activation and functional connectivity analyses as ROIs. Then using a GLM with covariates, we analyzed the preprocessed EPI data of the linguistic assay picture naming task within these ROIs. Specifically, every trial was modeled from the onset of the picture with a duration of 2 sec. For every participant, the correct and incorrect trials were coded as separate predictors to increase statistical power. Each correct trial was modeled by two covariates of interest (i.e., concept familiarity averaged from subject ratings and log-transformed word frequency). Furthermore, six head motion parameters as nuisance covariates were added to the model. All predictors and covariates of interest were convolved with the canonical HRF. We computed the main effects of concept familiarity and lexical frequency versus baseline (jitter trials) for voxels within the ROIs for each participant. Based on Graves et al. (2007) and Wilson et al. (2009), we expected that as concept familiarity/lexical frequency increases, the BOLD signal in the conceptual/lexical regions decreases. Therefore, if the averaged main effect of concept familiarity/lexical frequency across voxels of an ROI was significantly lower than 0, which indexes a negative correlation between brain activation and concept familiarity/lexical frequency, it suggests that this ROI is involved in conceptual/lexical processing during speech production. This analysis not only allows us to specify the cognitive role of an ROI, but also to infer the cognitive role of the functional connectivity between regions.

#### 6. Brain-Behavior Individual Differences Analysis

To understand the relationship between brain changes and behavioral effects (priming and interference) we correlated individual naming performance (using RTs but not accuracies, because we did not observe significant effects of relatedness on accuracies in either lag condition) with brain responses (ROI based whole-brain activation and functional connectivity analyses). To assess the neural priming effect, for each participant we extracted the averaged beta values of voxels in clusters identified in the univariate activation and functional connectivity analyses for the lag0 condition (lag0_related -lag0_unrelated). Then, across participants we correlated these neural priming effects with the magnitude of behavioral priming obtained in the RT analysis (lag0: naming RTs (ms) related – unrelated). For interference during naming, we extracted the averaged beta values of voxels in clusters identified in the voxel-wise and functional connectivity analyses for the lag2 condition (lag2_related -lag2_unrelated) and correlated them with the magnitude of behavioral interference found in the RT analysis (lag2: naming RTs (ms) related – unrelated). Because Pearson correlation can be highly susceptible to the effects of influential bivariate outliers (Rousselet and Pernet 2012), we conducted robust correlation analyses using Spearman’s ρ after removing outliers based on the bootstrapped Mahalanobis distance (Shepherd’s π test; Schwarzkopf et al. 2012). To limit the influence of data heteroscedasticity, correlations were considered significant if the Bonferroni corrected bootstrapped 95% CI did not include zero (Rousselet and Pernet 2012; Schwarzkopf et al. 2012).

## Results

### Behavioral Data

#### Priming-Interference Picture Naming Task

5.6% of RTs were removed from the RT analyses as a result of incorrect responses. Figure 2 shows mean RTs across the semantically related/unrelated and lag0/lag2 conditions with 95% confidence intervals (CI). The planned t-tests revealed priming (priming effect = 13 ms, 95% CI: 4-21 ms) in lag0 (t1(25) = 3.21, p =.004; t2(51) = 1.59, p =.12), and interference (interference effect = 24 ms, 95% CI: 14-34 ms) in lag2 (t1(25) = 4.94, p <.001; t2(51) = 2.98, p =.004). The error analysis did not reveal any difference in naming semantically related vs. unrelated targets in either lag0 or lag2 condition (*t*’s < 1). These results replicated the findings in a different group of participants from our previous study (Wei and Schnur 2019), demonstrating that past naming experience exerts a different influence on future naming depending on the interval between two naming occurrences.

**Figure 2.**
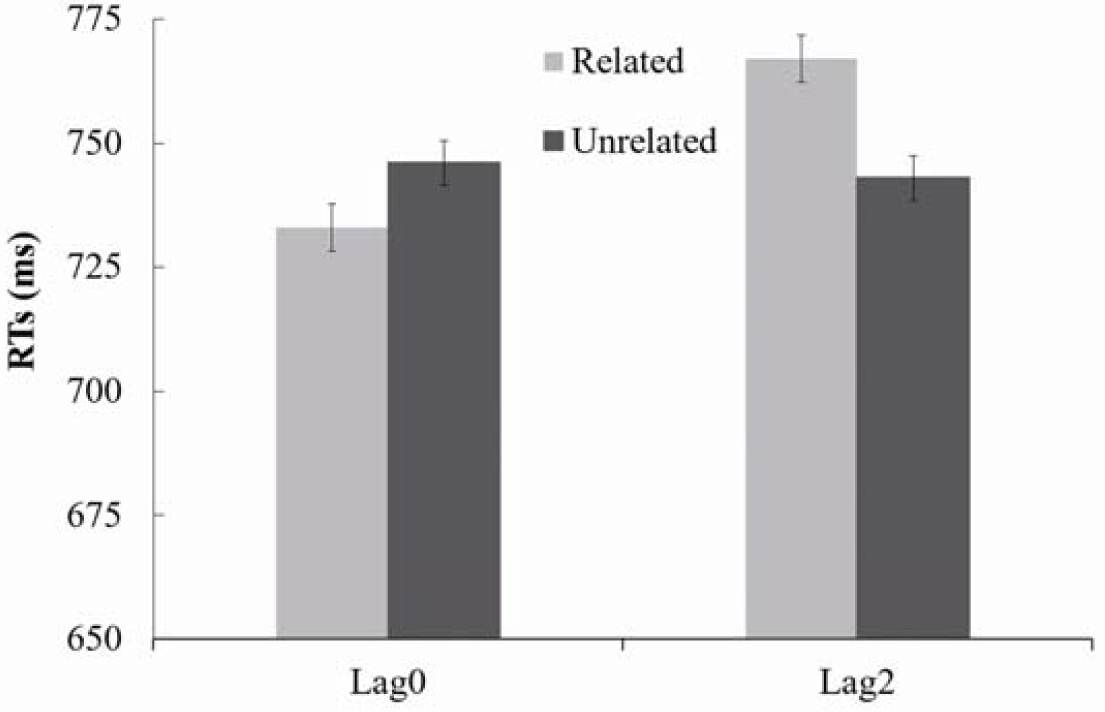
Mean RTs for naming target pictures in different conditions in the priming-interference picture naming task. The error bars represent 95% confidence intervals.

#### Linguistic Assay Picture Naming Task

Table 1 summarizes the distribution of concept familiarity, lexical frequency, RTs and accuracies associated with the 166 items used in the linguistic assay picture naming task. The distribution across items demonstrates significant variability across all dependent variables suggesting suitability of their use in correlational analyses. The distribution across participants suggests that all participants performed well in this naming task. Specifically, the mean naming accuracy across participants was 91% (SD = 5%; range 80 – 97%), and the mean RT across participants was 822 ms (SD = 82 ms; range 675 – 980 ms). Under the assumption that concept familiarity and lexical frequency respectively reflect conceptual and lexical processing during naming, they should correlate with naming performance (both RT and accuracy). To verify the contribution of concept familiarity and lexical frequency to naming, we performed pairwise correlations between concept familiarity, lexical frequency, and naming performance (RT and accuracy). As shown in Figure 3, all pairwise correlations were significant (*p*’s <.05) except the correlation between lexical frequency and concept familiarity (*p* =.36). Items with higher accuracy were named more quickly (demonstrating a lack of speed-accuracy trade-off). Importantly, both naming accuracy and RT were predicted by the concept familiarity and lexical frequency associated with pictures. Less familiar pictures as well as less frequent pictures were named more slowly with less accuracy. Because estimates of concept familiarity and lexical frequency did not correlate with each other, this suggests that these factors contributed independently to naming performance.

**Table 1.**
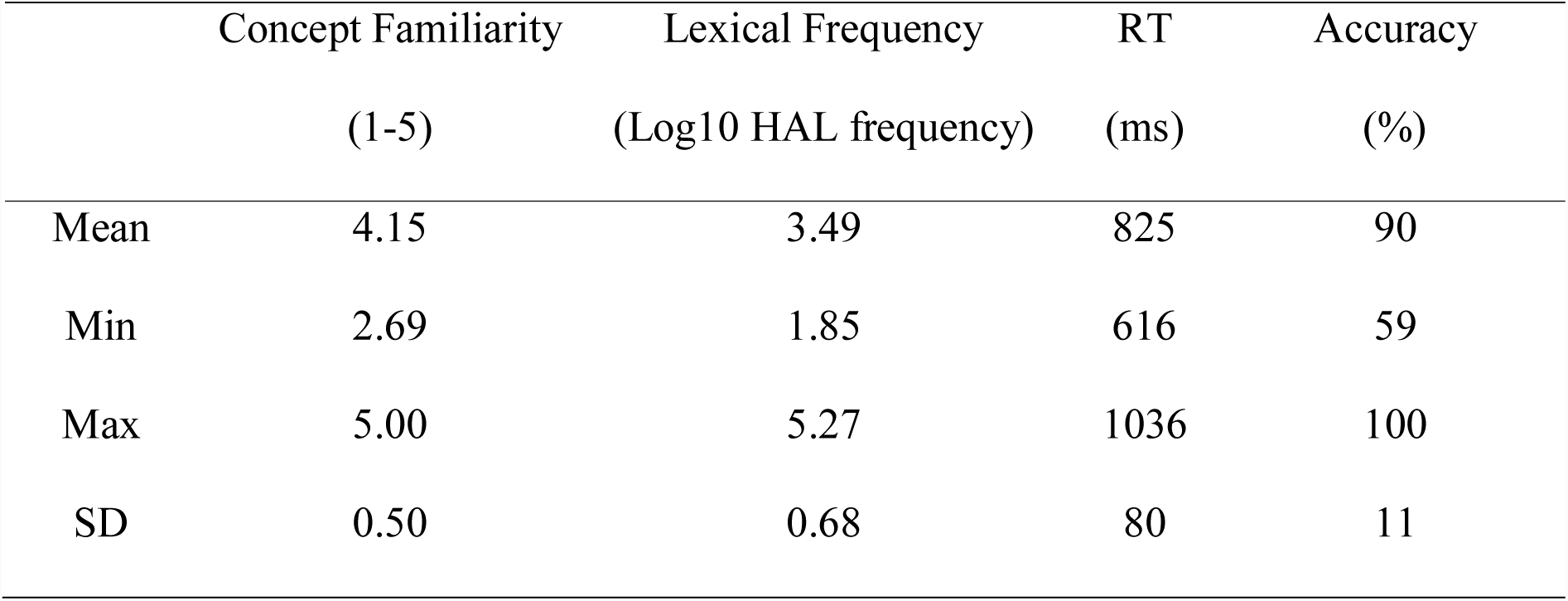
Characteristics of the stimuli used in the linguistic assay naming task. Variables associated with the 166 pictures, with mean, range and standard deviation calculated across items.

|  | Concept Familiarity | Lexical Frequency | RT | Accuracy |
| --- | --- | --- | --- | --- |
|  | (1-5) | (Log10 HAL frequency) | (ms) | (%) |
| Mean | 4.15 | 3.49 | 825 | 90 |
| Min | 2.69 | 1.85 | 616 | 59 |
| Max | 5.00 | 5.27 | 1036 | 100 |
| SD | 0.50 | 0.68 | 80 | 11 |

**Figure 3.**
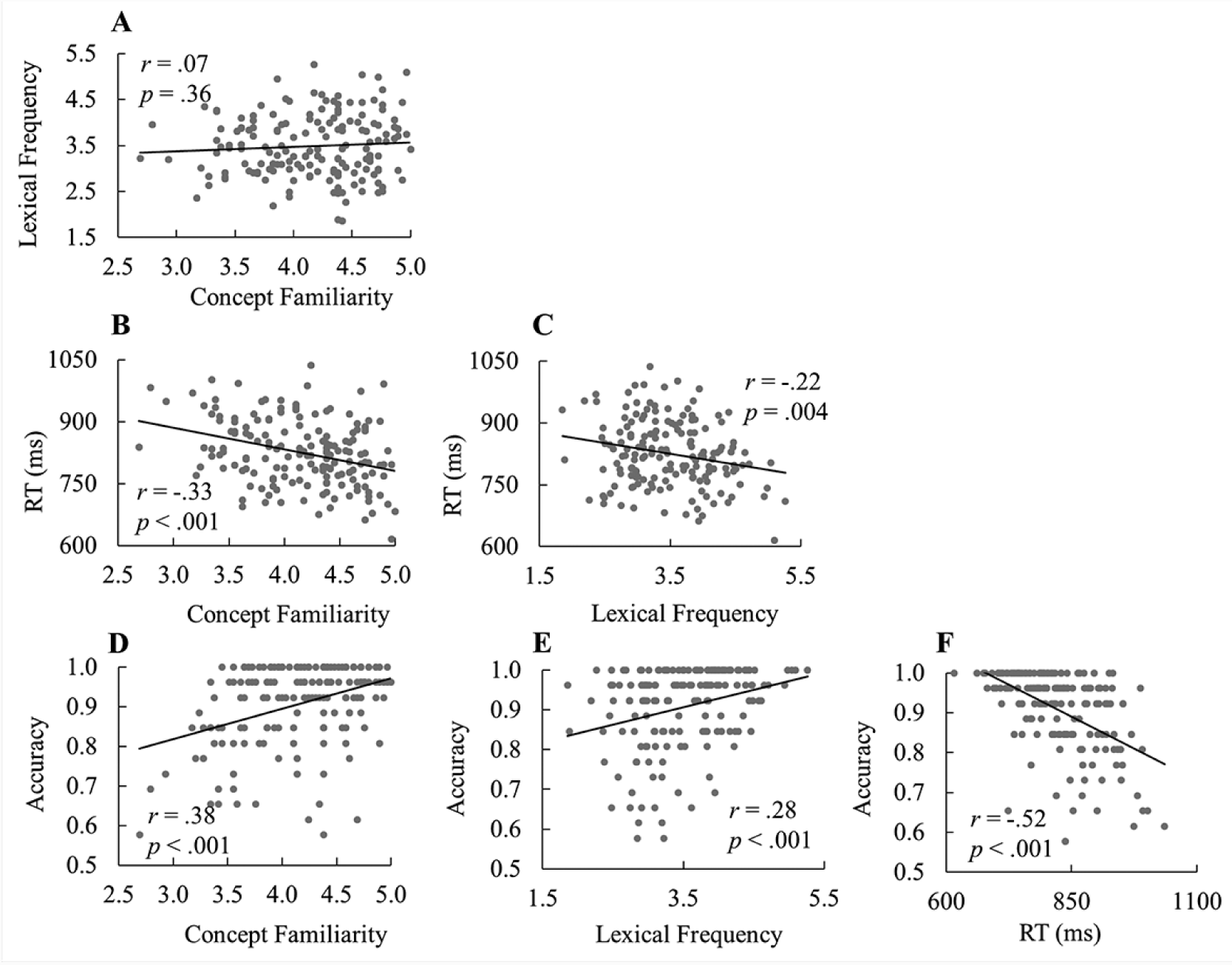
Pairwise correlations between the four behavioral variables in the linguistic assay picture naming task: concept familiarity and lexical frequency (A), RT and concept familiarity (B), RT and lexical frequency (C), accuracy and concept familiarity (D), accuracy and lexical frequency (E), accuracy and RT (F). Pearson correlation coefficients (r) and significance values (p) are shown. All correlations were significant except for that between concept familiarity and lexical frequency.

### Imaging Data

#### Brain Regions Modulated by Prior Naming Experience and their Cognitive Role in Word Production

In the fMRI analysis, we first identified brain regions that showed differential activation in response to semantically related vs. unrelated targets in lag0 and lag2 respectively (see Table 2 and Figure 4). For lag0 where the prime and target pictures were named adjacently and priming was observed behaviorally, semantically related vs. unrelated target pictures elicited greater activation in the bilateral ventral occipitotemporal cortex (vOTC). That bilateral vOTC activation is associated with priming during naming is consistent with previous studies of priming using different tasks (e.g., van Turennout et al. 2000; Chao et al. 2002; Wheatley et al. 2005; Gold et al. 2006).

**Table 2.**
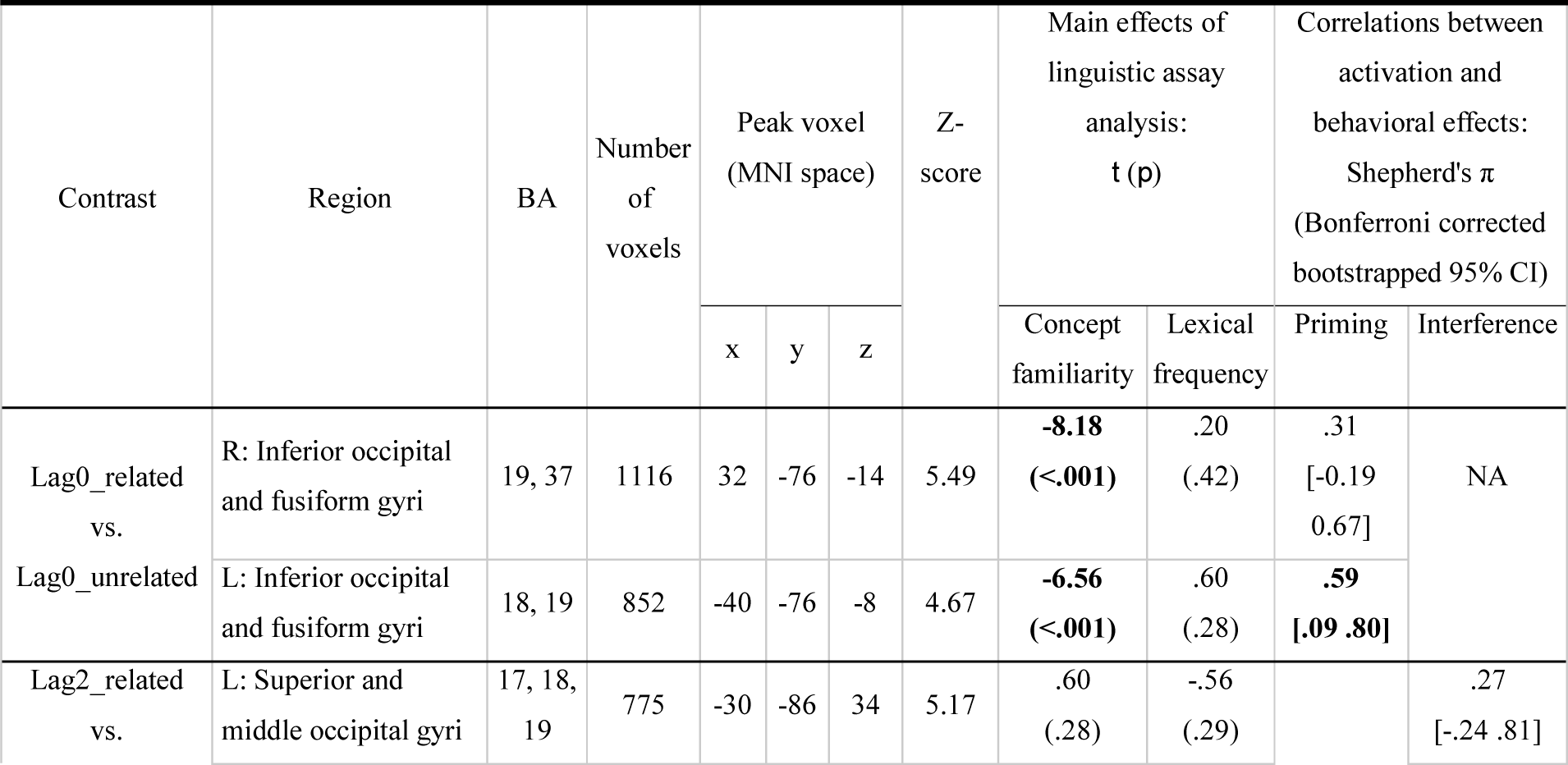
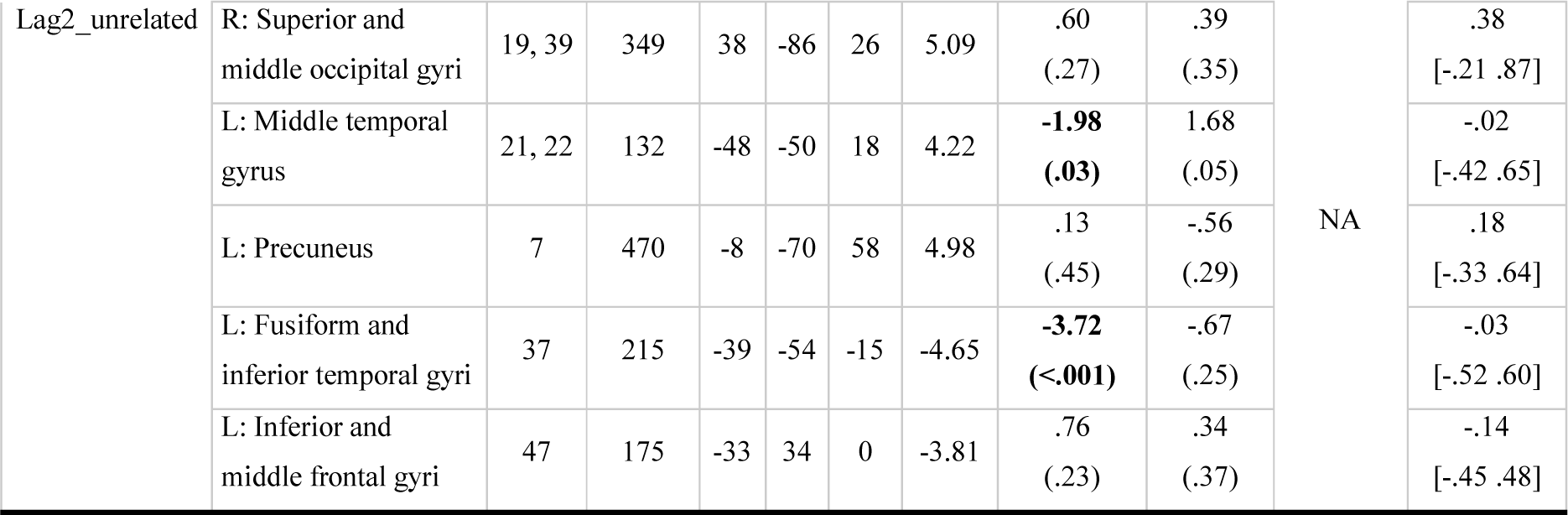
Brain regions demonstrating differential BOLD response for semantically related vs. unrelated targets for lag0 and lag2 (uncorrected voxel-wise *p* < 0.001 and cluster-level FWE corrected *p* < 0.05), region sensitivity to concept familiarity and lexical frequency, and across participant correlations between individual degree of region activation and participants’ RT magnitude of behavioral semantic priming/interference.

| Contrast | Region | BA | Number of voxels | Peak voxel (MNI space) | | | Z-score | Main effects of linguistic assay analysis: $t(p)$ | | Correlations between activation and behavioral effects: Shepherd's $\pi$ (Bonferroni corrected bootstrapped 95% CI) | |
| --- | --- | --- | --- | --- | --- | --- | --- | --- | --- | --- | --- |
|  |  |  |  | x | y | z |  | Concept familiarity | Lexical frequency | Priming | Interference |
| Lag0_related vs. Lag0_unrelated | R: Inferior occipital and fusiform gyri | 19, 37 | 1116 | 32 | -76 | -14 | 5.49 | **-8.18**<br>(**<.001**) | .20<br>(.42) | .31<br>[-0.19<br>0.67] | NA |
|  | L: Inferior occipital and fusiform gyri | 18, 19 | 852 | -40 | -76 | -8 | 4.67 | **-6.56**<br>(**<.001**) | .60<br>(.28) | **.59**<br>[.09 .80] |  |
| Lag2_related vs. | L: Superior and middle occipital gyri | 17, 18, 19 | 775 | -30 | -86 | 34 | 5.17 | .60<br>(.28) | -.56<br>(.29) |  | .27<br>[-.24 .81] |
| Lag2_unrelated | R: Superior and middle occipital gyri | 19, 39 | 349 | 38 | -86 | 26 | 5.09 | .60<br>(.27) | .39<br>(.35) | NA | .38<br>[-.21 .87] |
|  | L: Middle temporal gyrus | 21, 22 | 132 | -48 | -50 | 18 | 4.22 | **-1.98**<br>**(.03)** | 1.68<br>(.05) |  | -.02<br>[-.42 .65] |
|  | L: Precuneus | 7 | 470 | -8 | -70 | 58 | 4.98 | .13<br>(.45) | -.56<br>(.29) |  | .18<br>[-.33 .64] |
|  | L: Fusiform and inferior temporal gyri | 37 | 215 | -39 | -54 | -15 | -4.65 | **-3.72**<br>**(<.001)** | -.67<br>(.25) |  | -.03<br>[-.52 .60] |
|  | L: Inferior and middle frontal gyri | 47 | 175 | -33 | 34 | 0 | -3.81 | .76<br>(.23) | .34<br>(.37) |  | -.14<br>[-.45 .48] |

**Figure 4.**
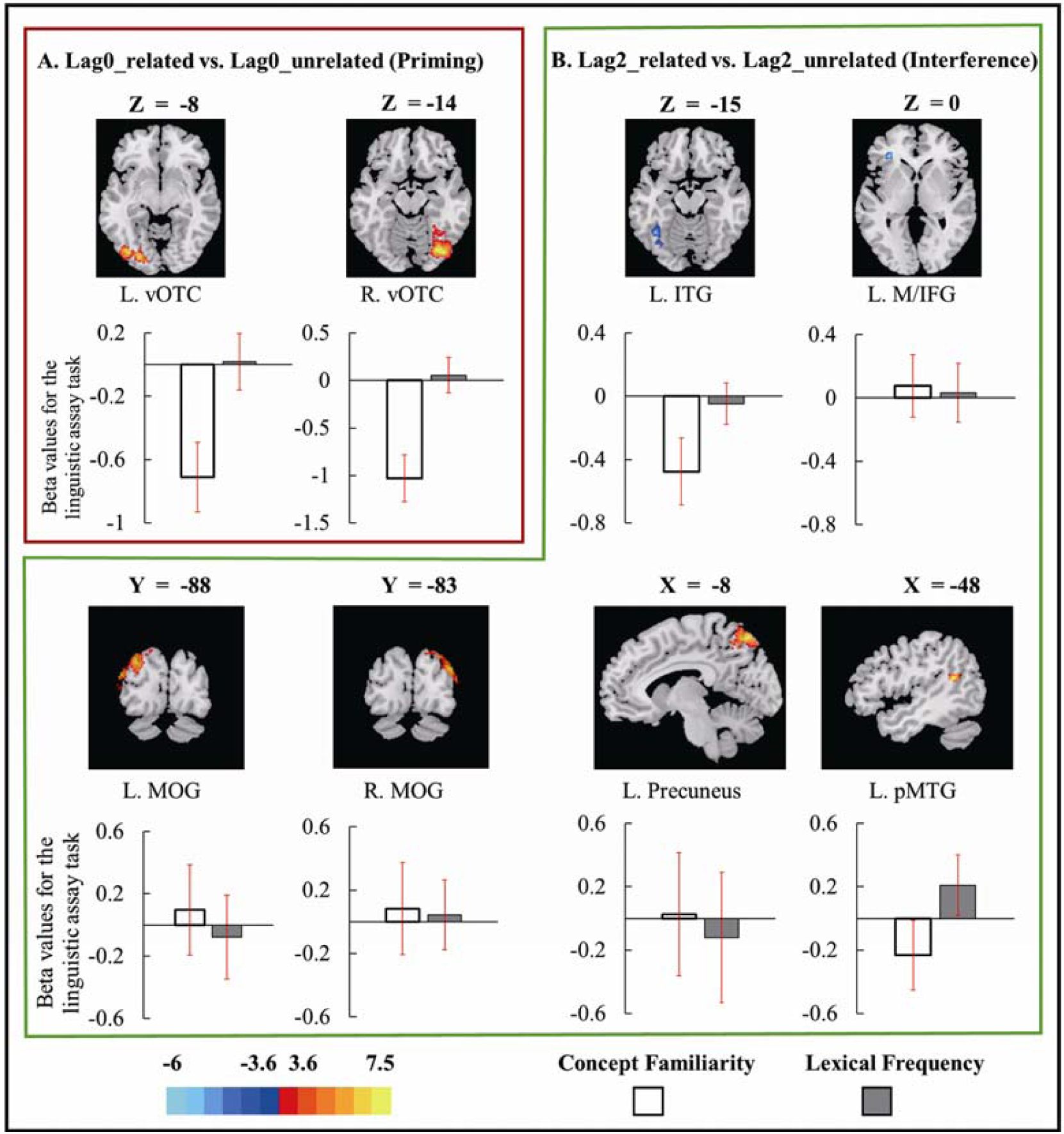
Brain regions showing differential BOLD responses during the naming of targets preceded by semantically related vs. unrelated primes for lag0 (A) and lag2 (B) respectively. Also depicted is brain region sensitivity to concept familiarity and lexical frequency. Red scale depicts the related > unrelated condition and the blue scale depicts unrelated > related targets. Averaged beta values associated with concept familiarity and lexical frequency are depicted in the bar plots. Estimates were extracted from the linguistic assay parametric analyses of concept familiarity and lexical frequency and averaged across the entire corresponding cluster listed in Table 2. Error bars represent 95% confidence interval of the mean across participants. L: left; R: right; ITG: inferior temporal gyrus; M/IFG: middle/inferior frontal gyrus; pMTG: posterior middle temporal gyrus; S/MOG: superior and middle occipital gyrus; VOTC: ventral occipitotemporal cortex.

Regarding interference in naming, for lag2 where the prime and target pictures were interleaved by two unrelated fillers, the left posterior middle temporal gyrus (pMTG) had greater activation for the semantically related than unrelated targets. That the left pMTG is associated with interference in naming is also found in other types of naming paradigms which create interference during naming (i.e. conceptual blocked naming, Schnur et al. 2009; de Zubicaray et al. 2013). In addition to the left pMTG, the bilateral superior/middle occipital gyrus (S/MOG) and precuneus also showed greater activation for semantically related vs. unrelated targets at lag2. The left inferior temporal gyrus (ITG) and middle/inferior frontal gyrus (M/IFG) were also involved in the conceptual effect but showed the opposite pattern (lag2-related < lag2-unrelated). To investigate whether priming and interference involve the same regions, we conducted a conjunction analysis (Nichols et al. 2005) by overlaying the map in lag0 (Figure 4A) and lag2 (Figure 4B). The result did not reveal any region associated with both effects, suggesting that different regions are involved in priming and interference.

Concept familiarity and lexical frequency are assumed to reflect conceptual and lexical level processing respectively and have been used to identify brain regions associated with conceptual and lexical processing (Graves et al. 2007; Wilson et al. 2009). Thus, to examine the cognitive role (conceptual vs. lexical) of brain regions associated with priming and interference, we tested these regions’ sensitivity to concept familiarity and lexical frequency in an independent picture naming fMRI experiment (i.e., the linguistic assay task). Following previous studies (Graves et al. 2007; Wilson et al. 2009), we only considered brain regions whose activation was negatively correlated with concept familiarity and lexical frequency. For lag0 (associated with priming), the bilateral vOTC which revealed differential activation when naming targets preceded by previously named semantically related vs. unrelated pictures, showed a significant effect of concept familiarity. This is consistent with a conceptual locus account of priming in naming (e.g., Lupker, 1988; Damian and Als 2005). For lag2 (associated with interference), among the brain regions whose activation was different between the related vs. unrelated targets (Figure 4B), the left ITG and pMTG revealed a significant effect of concept familiarity, which suggests an association between interference and conceptual processing (e.g., Biegler et al. 2008; Campanella and Shallice 2011; Wei and Schnur 2016). Other regions neither showed a significant effect of concept familiarity nor lexical frequency, suggesting that these regions are either involved in the priming or interference effect, but at a different level of the language production process, or are part of the conceptual and/or lexical process not captured by concept familiarity or lexical frequency. See details of statistical results in Table 2.

#### Functional Connectivity Modulated by Prior Naming Experience and its Cognitive Role in Word Production

Using the significant clusters identified in the activation analysis as seed regions, we conducted gPPI analyses to investigate how functional connectivity associated with these seed regions was modulated by relatedness in lag0 and lag2 respectively. The results are summarized in Table 3. For the priming effect when comparing the related vs. unrelated targets of lag0, we found stronger functional connectivity between the left vOTC seed region and the left MOG (also see Figure 5A). For the same contrast for the interference effect at lag2, the left ITG seed region showed stronger functional connectivity with bilateral lingual gyri, while left pMTG showed weaker functional connectivity (unrelated > related) with the left angular gyrus (AG).

**Table 3.** Brain regions showing differential functional connectivity with clusters identified in the whole-brain univariate activation analysis (see Table 2) for semantically related vs. unrelated targets of lag0 and lag2 respectively (uncorrected voxelwise *p* < 0.001 and cluster-level FWE corrected *p* < 0.050), their sensitivity to concept familiarity and lexical frequency, and correlations between functional connectivity strength and magnitude of RT semantic priming/interference. L: left; B: bilateral; VOTC: ventral occipitotemporal cortex; pMTG: posterior middle temporal gyrus; ITG: inferior temporal gyrus.

| | Seed | Connected region | BA | Number of voxels | Peak voxel (MNI space) | | | Z-score | Main effect of linguistic assay analysis: $t(p)$ | | Correlations between functional connectivity and behavior: $r(p)$ | |
| --- | --- | --- | --- | --- | --- | --- | --- | --- | --- | --- | --- | --- |
|  |  |  |  |  | x | y | z |  | Concept familiarity | Lexical frequency | Priming | Interference |
| Lag0_related vs. Lag0_unrelated | L. vOTC | L. Middle occipital gyrus | 19 | 488 | -28 | -78 | 14 | 4.62 | -1.29<br>(.10) | .51<br>(.30) | .08<br>[-.43 .44] | NA |
| Lag2_related vs. Lag2_unrelated | L. pMTG | L. Angular gyrus | 39 | 181 | -48 | -67 | 41 | -4.01 | -.81<br>(.21) | **-2.82**<br>**(.005)** | NA | **-.62**<br>**[-.93 -.19]** |
|  | L. ITG | B. Lingual gyrus | 18 | 407 | -12 | -80 | -4 | 5.07 | **-3.25**<br>**(.001)** | -.32<br>(.38) |  | .05<br>[-.46 .66] |

**Figure 5.**
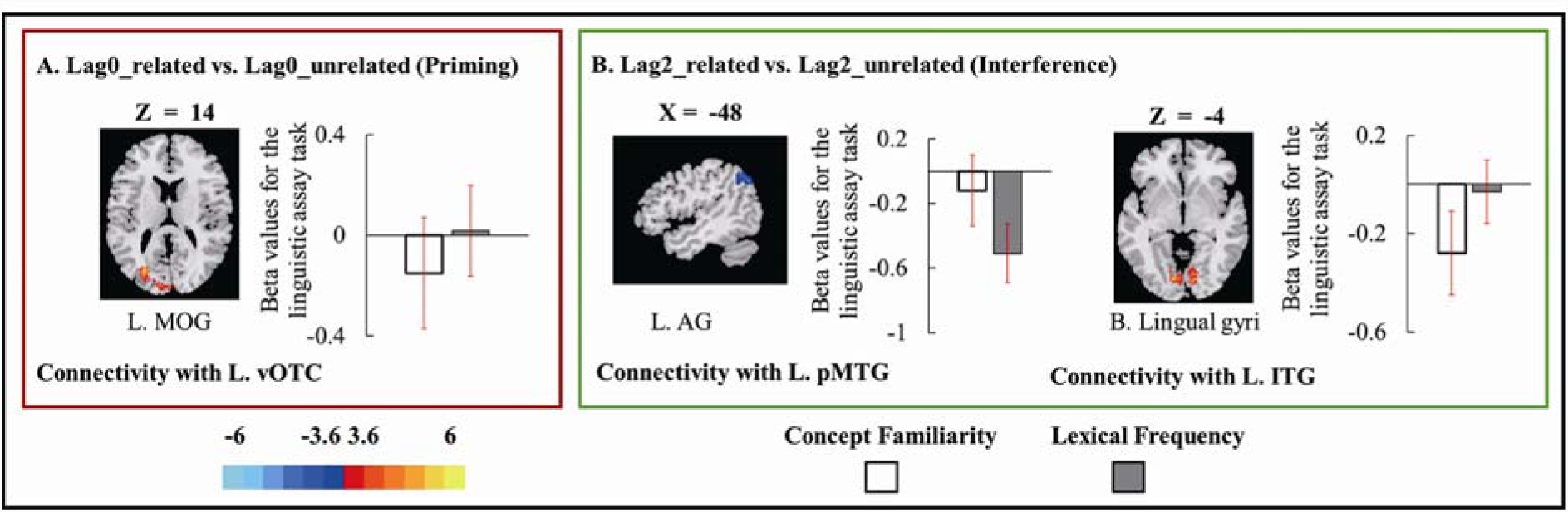
Brain regions showing differential functional connectivity during the naming of targets preceded by semantically related vs. unrelated primes for lag0 (A) and lag2 (B) respectively. Also depicted is brain region sensitivity to concept familiarity and lexical frequency. Red scale presents related > unrelated functional connectivity and blue scale presents unrelated > related functional connectivity. Averaged beta values associated with concept familiarity and lexical frequency are depicted in the bar plots. Estimates were extracted from linguistic assay parametric analyses of concept familiarity and lexical frequency and averaged across the entire corresponding cluster listed in Table 3. Error bars represent 95% confidence interval of the mean across participants. L: left; R: right; ITG: inferior temporal gyrus; pMTG: posterior middle temporal gyrus; S/MOG: superior and middle occipital gyrus; vOTC: ventral occipitotemporal cortex.

To investigate how the functional connectivity identified above contributes to priming and interference, we tested the cognitive role of the regions with which the connectivity associated. Using the same procedure as we did for the ROIs identified in the univariate activation analysis, we found that the left AG, whose functional connectivity strength with the left pMTG was affected by semantically related vs. unrelated targets in lag2, showed sensitivity to lexical frequency. Because the left pMTG was sensitive to concept familiarity (Figure 4B), we propose that the left pMTG-AG connection plays a role in mapping conceptual to lexical information.

#### Brain-Behavior Individual Differences Analysis: Relationships between Behavior and Brain Changes Due to Prior Naming Experience

To test whether brain activation and functional connectivity changes were related to the behavioral RT effects (priming and interference), we correlated the RT magnitude of participants’ priming and interference effects with their corresponding brain region activation of ROIs identified in the univariate activation analysis and connection strength from the functional connectivity analysis. For the two ROIs (Figure 4A) and one connection (Figure 5A) associated with priming, we found that the RT magnitude of priming was positively correlated with the univariate activation difference between related vs. unrelated targets in the left vOTC ROI (π =.59, bootstrapped 98.33% CI (95% CI adjusted for 3 comparisons) [.09,.80], see Figure 6A for the scatterplot). For the six ROIs (Figure 4B) and two connections (Figure 5B) associated with interference, only the connectivity strength of left pMTG-AG connection was negatively correlated with the RT magnitude of interference (π = -.62, bootstrapped 99.38% CI (95% CI adjusted for 8 comparisons) [-.93 -.19], see Figure 6B for the scatterplot). See Tables 2 and 3 for statistical details. All scatterplots are shown in Supplementary Materials Figure 1.

**Figure 6.**
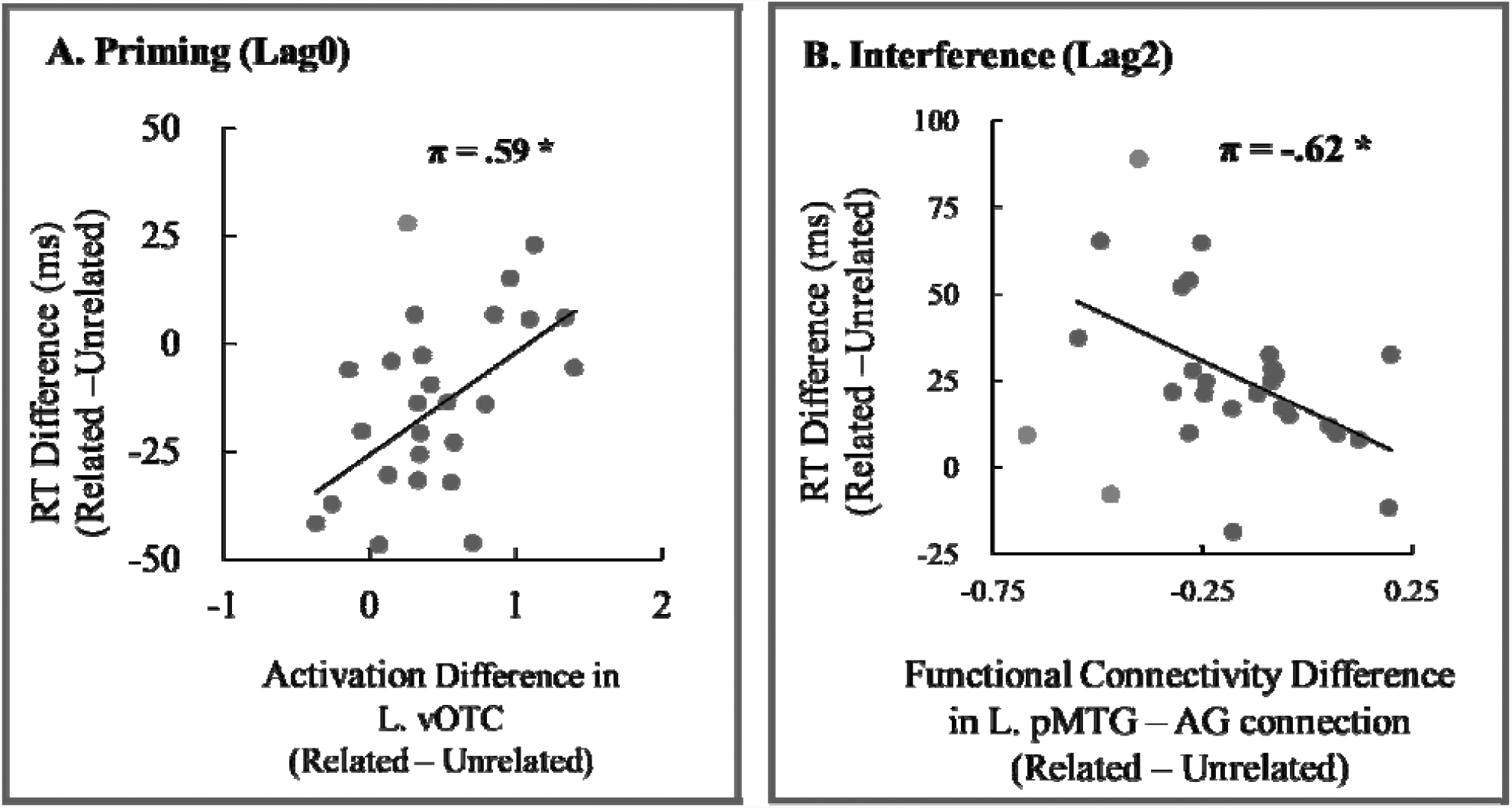
Neural signatures associated with priming and interference. Panel A depicts the significant positive correlation between the magnitude of the RT difference (related vs. unrelated) at lag 0 (priming; the larger the negative difference, the larger the priming) and the magnitude of the activation difference (related – unrelated) in the left vOTC. Panel B depicts the significant negative correlation between the magnitude of the RT difference (related vs. unrelated) at lag 2 (interference; the larger the positive difference, the larger the interference) and the magnitude of the connection strength difference (related – unrelated; interference) between the left pMTG and left AG. Detected outliers shown with gray dots. L: left; AG: angular gyrus; pMTG: posterior middle temporal gyrus; vOTC: ventral occipitotemporal cortex

## Discussion

Using fMRI, we addressed the neurological and cognitive mechanisms underlying how producing a word speeds up (priming) or slows down (interference) depending on prior naming experience (Wei and Schnur 2019). In a simple picture naming paradigm, participants named pictures one at a time in the scanner. We elicited priming and interference during the same task, naming, within the same participants and during the naming of the same items, depending on the recency of semantically related and unrelated previously named targets. We show that distinct brain regions and connectivity involved in conceptual and lexical stages during naming were associated with priming and interference effects respectively. Below we discuss how these key findings uncover the neural and cognitive basis of priming and interference due to previous speech and thus inform existing models of word production.

### Priming in naming is a conceptual level phenomenon

The first goal of this study was to provide neural evidence of whether priming in naming occurs at a conceptual level of the language production system (e.g., Lupker, 1988;), a lexical level (Navarrete et al. 2014), or is potentially not part of the language production process at all (Schnur et al. 2006; Oppenheim et al. 2010; Roelofs 2018). This scientific question is important because the access of conceptual representations during naming is usually not fully addressed in the computational modeling of word production (e.g., Dell et al. 1997; Howard et al. 2006; Oppenheim et al. 2010), even though conceptual representation access is part of the naming process (cf. Levelt et al. 1999) and when damaged leads to profound language problems (e.g., Warrington and Shallice 1984; for a review see Patterson et al. 2007). Behaviorally we observed that when prime and target pictures were named one after the other (lag0), semantically related targets were named faster than unrelated targets. Neurally, naming semantically related vs. unrelated primes elicited greater activation in the bilateral vOTC when naming a subsequent target and this change in activation was related to the degree of an item’s conceptual familiarity, a measure of conceptual knowledge. Further, the left vOTC activation difference was directly tied to naming behavior, as the left vOTC activation difference was positively correlated with the RT difference (priming) between the related and unrelated condition. Lastly, when participants named semantically related vs. unrelated targets, we found stronger functional connectivity between the left vOTC and the left MOG, a region for early visual processing, but the degree of functional connectivity was not significantly related to the RT priming effect or our linguistic assay variables (conceptual familiarity and lexical frequency). We interpret these results to suggest that first, because of the direct relationship between left vOTC activity and both magnitude of priming during naming and an item’s degree of conceptual familiarity, it is the left vOTC which subserves the conceptual representations that are modulated by past naming experience to produce faster naming. Second, although the bilateral vOTC and connections with the left MOG are involved when naming is primed by previously naming semantically related pictures, the lack of relationship with naming RTs and an item’s linguistic variables suggest these regions are involved at a different stage of picture naming other than conceptual and lexical stages. Given the cognitive role of the left vOTC and MOG, we speculate that this connection reflects the mapping between low level visual and conceptual information.

Although to our knowledge no study has examined the neural basis of priming in naming, the association between the vOTC and priming is demonstrated in other tasks, such as word reading (e.g., Wheatley et al. 2005) and lexical decision (e.g., Gold et al. 2006; Kuperberg et al. 2008). The involvement of vOTC in priming during word reading is thought to reflect spreading activation between semantically related representations in the conceptual system (Wheatley et al. 2005; Gold et al. 2006). Thus, our findings provide strong evidence that priming during naming occurs at a conceptual level of processing in the language system in the left vOTC.

### Interference during naming occurs at multiple stages of language production

We demonstrate for the first time to our knowledge, that in addition to a change in regional brain activation, a change in functional connectivity strength between brain regions is a contributory neural mechanism for interference in naming. Replicating our previous work (Wei and Schnur 2019), when prime and target pictures were interleaved by two unrelated pictures (lag2), participants were slower to name semantically related compared to unrelated targets.

Similar to previous studies focusing on the neural basis of interference in naming, we observed that naming semantically related vs. unrelated primes evoked differential brain activation in the left temporal lobe when naming targets, i.e., left pMTG (de Zubicaray et al. 2006; 2014; Schnur et al. 2009) and left ITG (de Zubicaray et al. 2006; 2013; 2014), along with other regions in the parietal (precuneus cortex), frontal (M/IFG), and occipital (MOG) lobes. Importantly, we found that functional connectivity associated with specific regions was modulated by semantically related vs. unrelated primes. After naming semantically related vs. unrelated primes, functional connectivity between the left ITG and bilateral lingual gyri was increased and functional connectivity between the left pMTG and left AG was decreased. The contribution of the change in functional connectivity strength to interference was further confirmed by its correlation with the RT magnitude of interference. We found that the stronger the functional connectivity between the left pMTG and left AG, the smaller the interference effect across individuals, a pattern not observed between the left ITG and bilateral lingual gyri. This result suggests that prior naming experience changes the efficiency of communication between left pMTG and AG which affects future naming performance, creating interference.

How are these regions and connection dynamics contributing to how interference in naming occurs? The involvement of the temporoparietal junction (TPJ) for interference in naming has been taken as evidence that interference is a lexical effect (de Zubicarary et al. 2014; 2015), as this region is found elsewhere to be involved in lexical processing (Hickok and Poeppel 2007; Indefrey and Levelt 2004). However, as discussed in the Introduction, this region is also reported to be involved in conceptual processing (Martin 2007; Binder and Desai 2011). Our results tease apart this seemingly conflicting evidence. We observed two ROIs associated with the interference effect within the TPJ which we directly tested for their respective cognitive roles (conceptual vs. lexical) in an independent linguistic assay task. The first ROI was the left pMTG whose activation was modulated by semantically related vs. unrelated primes in lag2 (interference). We observed that activation in left pMTG was sensitive to concept familiarity, suggesting that this region was involved in conceptual processing. The second ROI associated with interference was the left AG, whose functional connectivity with the left pMTG was modulated by semantically related vs. unrelated primes in lag2. The activation in the left AG ROI was found to be sensitive to lexical frequency, indicating its role in lexical processing (Booth et al. 2003; Wilson et al. 2009). Most importantly, the connectivity strength between the left pMTG and AG was correlated with magnitude of interference. Thus, our results suggest that interference in naming occurs with changes in conceptual processing and coupling between conceptual and lexical processes.

Lastly, we found that interference was associated with an additional region, the left ITG and its functional connectivity with bilateral lingual gyri. Activation within the left ITG and bilateral lingual gyri was sensitive to concept familiarity, suggesting that both regions implicated in interference were involved at a conceptual level of language processing. However, neither activation of the left ITG nor its connectivity with bilateral lingual gyri changed with the magnitude of interference in naming. This suggests that these regions and connection are involved in some unidentified aspect of interference during naming. Future studies are needed to understand the specific roles in word production.

### Implications for computational modeling of language production

The profile of the findings from this experiment provides insight for computational models of word production. First, no existing computational models of word production currently account for the mechanisms that we identified for interference in naming. Three computational models of word production explain how the neurotypical language system gives rise to the behavioral changes caused by prior naming experience. In the Howard et al. (2006) and Oppenheim et al. (2010) models, prior naming experience changes two levels in naming: the strength of connections between conceptual and lexical representations, which further affects activation levels of lexical representations. The Roelofs model (2018) assumes changes in both conceptual and lexical levels. Prior naming experience affects the activation of conceptual representations. This activation passes down to the lexical level and induces differential activation of lexical representations. In this study, we showed that interference emerged along with changes at two levels: 1) the conceptual level of processing (the left ITG and left pMTG); and 2) the connection between conceptual and lexical processes (functional connectivity between the left pMTG and left AG). Different from all three models, we did not observe that activation in a lexical region, the left AG, was modulated by prior naming experience. To see whether this failure to detect the activation change at the lexical level of processing was due to relative conservative threshold in the whole brain activation analysis, we did a posthoc analysis by defining the left AG found in the functional connectivity analysis as an ROI and compared the activation of this ROI between the related vs. unrelated condition in lag2. This posthoc analysis showed that this left AG ROI had greater activation in the related vs. unrelated condition (t(25) = 2.09, p =.047). This result is tentative and it leaves open the possibility there is no lexical level effect of interference. Together, our results suggest that the computational models of word production consider how changes at different levels work together to induce interference in naming.

Second, to fully capture naming performance, computational models of naming should refine how the conceptual level of processing is affected by prior naming experience and induces the priming effect. None of the three models explicitly simulate the priming effect caused by naming semantically related primes. This is mainly because some of these models (Oppenheim et al. 2010; Roelofs 2018) assume that the priming effect in naming is due to expectation rather than changes within the language system, although most studies observing priming in naming assume that it occurs at the conceptual level of processing during naming (Huttenlocher and Kubicek 1983; Lupker 1988; Biggs and Marmurek 1990; Sperber et al. 1979). Elsewhere we demonstrated that priming in naming was still evident when the influence of expectation was minimized (Wei and Schnur 2019). Here, using the same paradigm we found that priming was associated with vOTC, a region involved in conceptual processing, while no regions associated with expectation (Gold et al. 2006) were identified. This result confirms a conceptual locus of priming in naming and a successful computational model of word production should account for both priming and interference. In addition, we found that priming and interference in naming were associated with conceptual processing in different brain regions. This suggests that the paradoxical effects arise due to modulations of different aspects of the conceptual system (Wei and Schnur 2016). It is not clear how current models of word production with a simplified conceptual system (unified conceptual nodes, Howard et al. 2006; Roelofs 2018 or distributed features, Oppenheim et al. 2010) account for both effects at the conceptual level of processing.

In conclusion, this study elucidated the neural and cognitive mechanisms underlying the paradoxical effects of priming and interference due to prior naming experience by measuring both brain activation and functional connectivity profiles while examining their cognitive role in naming. We demonstrated that immediate prior naming events change the engagement of conceptual regions for future naming resulting in a speed up in naming (priming), while distal naming events engage both conceptual and lexical regions and connections between them creating a slowing down in naming, i.e. interference. Our findings provide direct implications for computational models of word production.

## Acknowledgements

We thank Bowie Lin for helping extract RTs of naming experiments. We presented results at the Society for the Neurobiology of Language (Baltimore, MD 2017).

## Address of the corresponding author

Tatiana T. Schnur, Ph.D.

Baylor College of Medicine, Department of Neurosurgery, BCM 240

One Baylor Plaza, Houston, TX 77030

## References

Ashburner J. 2007. A fast diffeomorphic image registration algorithm. Neuroimage. 38(1): 95–113.

Balota DA, Yap MJ, Hutchison KA, Cortese MJ, Kessler B, Loftis B. Treiman R. 2007. The English lexicon project. Behavior research methods. 39: 445–459.

Belke E. 2013. Long-lasting inhibitory semantic context effects on object naming are necessarily conceptually mediated: Implications for models of lexical-semantic encoding. Journal of Memory & Language. 69: 228–256.

Biegler KA, Crowther JE, Martin RC. 2008. Consequences of an inhibition deficit for word production and comprehension: Evidence from the semantic blocking paradigm. Cognitive Neuropsychology. 25: 493–527.

Biggs TC, Marmurek HH. 1990. Picture and word naming: Is facilitation due to processing overlap?. The American Journal of Psychology. 103: 81–100.

Binder JR, Desai RH. 2011. The neurobiology of semantic memory. Trends in Cognitive Sciences. 15: 527–536.

Booth JR, Burman DD, Meyer JR, Gitelman DR, Parrish TB, Mesulam MM. 2003. Relation between brain activation and lexical performance. Human brain mapping. 19(3): 155–169.

Brodeur MB, Guérard K, Bouras M. 2014. Bank of standardized stimuli (BOSS) phase II: 930 new normative photos. PloS One. 9 e106953.

Brown AS. 1981. Inhibition in cued retrieval. Journal of Experimental Psychology: Human Learning and Memory. 7(3): 204.

Campanella F, Shallice T. 2011. Refractoriness and the healthy brain: a behavioural study on semantic access. Cognition. 118: 417–431.

Caramazza A. 1997. How many levels of processing are there in lexical access?. Cognitive Neuropsychology. 14: 177–208.

Chao LL, Weisberg J, Martin A. 2002. Experience-dependent modulation of category-related cortical activity. Cerebral Cortex. 12:545–51

Damian MF, Vigliocco G, Levelt WJ. 2001. Effects of semantic context in the naming of pictures and words. Cognition. 81: B77–B86.

de Zubicaray G, Johnson K, Howard D, McMahon K. 2014. A perfusion fMRI investigation of thematic and categorical context effects in the spoken production of object names. Cortex. 54: 135–149.

de Zubicaray G, McMahon K, Howard D. 2015. Perfusion fMRI evidence for priming of shared feature-to-lexical connections during cumulative semantic interference in spoken word production. Language Cognition & Neuroscience. 30: 261–272.

de Zubicaray G, McMahon K, Eastburn M, Pringle A. 2006. Top-down influences on lexical selection during spoken word production: A 4T fMRI investigation of refractory effects in picture naming. Human Brain Mapping. 27: 864–873.

de Zubicaray GI, Hansen S, McMahon KL. 2013. Differential processing of thematic and categorical conceptual relations in spoken word production. Journal of Experimental Psychology: General. 142(1): 131–142.

Dell GS. 1986. A spreading-activation theory of retrieval in sentence production. Psychological Review. 93: 283–321.

Dell GS, Schwartz MF, Martin N, Saffran EM, Gagnon DA. 1997. Lexical access in aphasic and nonaphasic speakers. Psychological review. 104(4): 801–838.

Fiez JA, Tranel D. 1997. Standardized stimuli and procedures for investigating the retrieval of lexical and conceptual knowledge for actions. Memory & Cognition. 25: 543–569.

Finocchiaro C, Caramazza A. 2006. The production of pronominal clitics: Implications for theories of lexical access. Language & Cognitive Processes. 21: 141–180.

Funnell E, De Mornay Davies P. 1997. JBR: A reassessment of concept familiarity and a category-specific disorder for living things. Neurocase. 2: 461–474.

Gold BT, Balota DA, Jones SJ, Powell DK, Smith CD, Andersen AH. 2006. Dissociation of automatic and strategic lexical-semantics: functional magnetic resonance imaging evidence for differing roles of multiple frontotemporal regions. The Journal of Neuroscience. 26: 6523–6532.

Graves WW, Grabowski TJ, Mehta S, Gordon JK. 2007. A neural signature of phonological access: distinguishing the effects of word frequency from familiarity and length in overt picture naming. Journal of Cognitive Neuroscience. 19: 617–631.

Hirsh KW, Funnell E. 1995. Those old familiar things: Age of acquisition familiarity and lexical access in progressive aphasia. Journal of Neurolinguistics. 9: 23–32.

Howard D, Nickels L, Coltheart M, Cole-Virtue J. 2006. Cumulative semantic inhibition in picture naming: Experimental and computational studies. Cognition. 100: 464–482.

Huttenlocher J, Kubicek LF. 1983. The source of relatedness effects on naming latency. Journal of Experimental Psychology: Learning Memory & Cognition. 9: 486–496.

Indefrey P, Levelt WJ. 2004. The spatial and temporal signatures of word production components. Cognition. 92: 101–144.

Kuperberg GR, Lakshmanan BM, Greve DN, West WC. 2008. Task and semantic relationship influence both the polarity and localization of hemodynamic modulation during lexico-semantic processing. Human brain mapping. 29(5): 544–561.

Lambon Ralph MA, Graham KS, Ellis W, Hodges JR. 1998. Naming in semantic dementia--what matters? Neuropsychologia. 36: 775–784.

Levelt WJ, Roelofs A, Meyer AS. 1999. A theory of lexical access in speech production. Behavioral & Brain Sciences. 22: 1–38.

Lupker SJ. 1988. Picture naming: An investigation of the nature of categorical priming. Journal of Experimental Psychology: Learning Memory & Cognition. 14: 444–455.

Martin A. 2007. The representation of object concepts in the brain. Annual Review of Psychology. 58: 25–45.

McClelland JL, Rogers TT. 2003. The parallel distributed processing approach to semantic cognition. Nature Reviews Neuroscience. 4: 310–322.

McRae K, Boisvert S. 1998. Automatic semantic similarity priming. Journal of Experimental Psychology: Learning Memory & Cognition. 24: 558–572.

Meyer DE, Schvaneveldt RW. 1971. Facilitation in recognizing pairs of words: evidence of a dependence between retrieval operations. Journal of Experimental Psychology. 90: 227–234.

Meyer DE, Schvaneveldt RW, Ruddy MG. 1975. Loci of contextual effects on visual word-recognition. In P. M. A. Rabbitt (Ed.) Attention and Performance V (pp. 98–118). London England: Academic Press.

Neely JH. 1991. Semantic priming effects in visual word recognition: A selective review of current findings and theories. Basic Processes in Reading: Visual Word Recognition. 11: 264–336.

Nichols T, Brett M, Andersson J, Wager T, Poline JB. 2005. Valid conjunction inference with the minimum statistic. Neuroimage. 25(3): 653–660.

Oppenheim G. M. Dell G. S. & Schwartz M. F. (2010). The dark side of incremental learning: A model of cumulative semantic interference during lexical access in speech production. Cognition 114 227–252.

Patterson K, Nestor PJ, Rogers TT. 2007. Where do you know what you know? The representation of semantic knowledge in the human brain. Nature Reviews Neuroscience. 8: 976–987.

Protopapas A. 2007. Check Vocal: A program to facilitate checking the accuracy and response time of vocal responses from DMDX. Behavior Research Methods. 39: 859–862.

Rastle K, Coltheart M. 1999. Lexical and nonlexical phonological priming in reading aloud. Journal of Experimental Psychology: Human Perception & Performance. 25: 461–481.

Roelofs A. 2018. A unified computational account of cumulative semantic, semantic blocking, and semantic distractor effects in picture naming. Cognition. 172: 59–72.

Rogers TT, Patterson K, Jefferies E, Lambon Ralph MA. 2015. Disorders of representation and control in semantic cognition: Effects of familiarity typicality and specificity. Neuropsychologia. 76: 220–239.

Rossell SL, Price CJ, Nobre AC. 2003. The anatomy and time course of semantic priming investigated by fMRI and ERPs. Neuropsychologia. 41(5): 550–564.

Rousselet GA, Pernet CR. 2012. Improving standards in brain-behavior correlation analyses. Frontiers in human neuroscience. 6: 119.

Schnur TT, Schwartz MF, Brecher A, Hodgson C. 2006. Semantic interference during blocked-cyclic naming: Evidence from aphasia. Journal of Memory & Language. 54: 199–227.

Schnur TT, Schwartz MF, Kimberg DY, Hirshorn E, Coslett HB, Thompson-Schill SL. 2009. Localizing interference during naming: convergent neuroimaging and neuropsychological evidence for the function of Broca’s area. Proceedings of the National Academy of Sciences. 106: 322–327.

Schwarzkopf DS, De Haas B, Rees G. 2012. Better ways to improve standards in brain-behavior correlation analysis. Frontiers in human neuroscience. 6: 200.

Strijkers K, Costa A, Thierry G. 2010. Tracking lexical access in speech production: electrophysiological correlates of word frequency and cognate effects. Cerebral Cortex. 20: 912–928.

Sperber RD, McCauley C, Ragain RD, Weil CM. 1979. Semantic priming effects on picture and word processing. Memory & Cognition. 7: 339–345.

Ulrich M, Adams SC, Kiefer M. 2014. Flexible establishment of functional brain networks supports attentional modulation of unconscious cognition. Human brain mapping. 35(11): 5500–5516.

van Turennout M, Ellmore T, Martin A. 2000. Long-lasting cortical plasticity in the object naming system. Nature neuroscience. 3(12): 1329–1334.

Vitkovitch M, Humphreys GW. 1991. Perseverant Responding in Speeded Naming of Pictures: It’s in the Links. Journal of Experimental Psychology. Learning Memory & Cognition. 17: 664–680.

Vitkovitch M, Cooper-Pye E, Leadbetter AG. 2006. Semantic priming over unrelated trials: Evidence for different effects in word and picture naming. Memory & Cognition. 34: 715–725.

Warrington EK, Shallice T. 1984. Category specific semantic impairments. Brain. 107(3): 829–853.

Wei, T., Schnur, TT. 2019. Being fast or slow at naming depends on recency of experience. Cognition. 182: 165–170.

Wei T, Schnur TT. 2016. Long-term interference at the semantic level: Evidence from blocked-cyclic picture matching. Journal of Experimental Psychology: Learning Memory & Cognition. 42: 149–157.

Wheatley T, Weisberg J, Beauchamp MS, Martin A. 2005. Automatic priming of semantically related words reduces activity in the fusiform gyrus. Journal of Cognitive Neuroscience. 17: 1871–1885.

Wheeldon LR, Monsell S. 1994. Inhibition of spoken word production by priming a semantic competitor. Journal of Memory & Language. 33: 332–356.

Wible CG, Han SD, Spencer MH, Kubicki M, Niznikiewicz MH, Jolesz FA,… Nestor P. 2006. Connectivity among semantic associates: an fMRI study of semantic priming. Brain and Language. 97(3): 294–305.

Wilson SM, Isenberg AL, Hickok G. 2009. Neural correlates of word production stages delineated by parametric modulation of psycholinguistic variables. Human Brain Mapping. 30: 3596–3608.

Woollams AM, Cooper-Pye E, Hodges JR, Patterson K. 2008. Anomia: A doubly typical signature of semantic dementia. Neuropsychologia. 46: 2503–2514.

