## Supplemental Figure 1 for "Neural and linguistic differences explain priming and interference during naming"

SM Figure 1. Correlations between behavior and brain changes due to prior naming experience. Panel A depicts correlations between the magnitude of the RT difference (related vs. unrelated) at lag 0 (priming; the larger the negative difference, the larger the priming) and the magnitude of the brain difference (related – unrelated) associated with priming. Panel B depicts the correlation between the magnitude of the RT difference (related vs. unrelated) at lag 2 (interference; the larger the positive difference, the larger the interference) and the magnitude of the brain difference (related – unrelated; interference) associated with interference. Detected outliers shown with gray dots. L: left; R: right; B: bilateral; AG: angular gyrus; ITG: inferior temporal gyrus; M/IFG: middle/inferior frontal gyrus; MOG: middle occipital gyrus; pMTG: posterior middle temporal gyrus; vOTC: ventral occipitotemporal cortex.
